# Decoding Spatial Tissue Architecture: A Scalable Bayesian Topic Model for Multiplexed Imaging Analysis

**DOI:** 10.1101/2024.10.08.617293

**Authors:** Xiyu Peng, James W. Smithy, Mohammad Yosofvand, Caroline E. Kostrzewa, MaryLena Bleile, Fiona D. Ehrich, Jasme Lee, Michael A. Postow, Margaret K. Callahan, Katherine S. Panageas, Ronglai Shen

**Author notes:** Contributing authors.

## Abstract

Recent progress in multiplexed tissue imaging is advancing the study of tumor microenvironments to enhance our understanding of treatment response and disease progression. Cellular neighborhood analysis is a popular computational approach for these complex image data. Despite its popularity, there are significant challenges, including high computational demands that limit feasibility for largescale applications and the lack of a principled strategy for integrative analysis across images. This absence hampers the precise and consistent identification of spatial features and tracking of their dynamics over disease progression. To overcome these challenges, we introduce *SpatialTopic*, a spatial topic model designed to decode high-level spatial architecture across multiplexed tissue images. *SpatialTopic* integrates both cell type and spatial information within a topic modelling framework, originally developed for natural language processing and adapted for computer vision. Spatial information is incorporated into the flexible design of documents, representing densely overlapping regions in images. We employ an efficient collapsed Gibbs sampling algorithm for model inference. We benchmarked the performance against five state-of-the-art algorithms through various case studies using different single-cell spatial transcriptomic and proteomic imaging platforms across different tissue types. We show that *SpatialTopic* is highly scalable on large-scale image datasets with millions of cells, along with high precision and interpretability. Our findings demonstrate that *SpatialTopic* consistently identifies biologically and clinically significant spatial “topics” such as tertiary lymphoid structures (TLSs) and tracks dynamic changes in spatial features over disease progression. Its computational efficiency and broad applicability across various molecular imaging platforms will enhance the analysis of large-scale tissue imaging datasets.

## Introduction

Recent advancements in multiplexed tissue imaging allow the profiling of RNA and protein expression in situ across thousands to millions of single cells within a whole-slide tissue context [1–5]. These technologies generate high-dimensional molecular imaging data, offering significant opportunities for a spatially resolved understanding of cellular heterogeneity and organization within tissues. Compared to other single-cell technologies (such as single-cell RNA-seq, flow cytometry), multiplexed imaging provides unique opportunities to examine spatial patterns of diverse cell types and characterize the tissue microenvironment of interest, which may play an essential role in understanding disease progression, tissue development, and mechanisms of treatment response [1, 2, 4–7]. One recent discovery in cancer, partly enabled by multiplexed spatially resolved omics data, is the presence of tertiary lymphoid structures (TLSs) in tumor tissues and its role in the adaptive antitumor immune response [8–11]. TLSs have been identified in a wide range of human cancers [9] and have demonstrated a promising positive association with improved outcomes in cancer patients who underwent immunotherapy [8].

While promising, the complex cellular architecture revealed by whole-slide multiplexed tissue imaging presents significant analytical challenges. Pathology images of tissue samples affected by certain diseases, such as cancer, are particularly complex, displaying abnormal cellular structures and significant variation between tumor samples. Currently, most analyses focus on individual images, examining elements such as cell densities and inter-cellular distances [1, 2, 6], or conducting basic spatial domain analyses that primarily focus on binarized tissue compartments, such as tumor versus stroma [12]. Associating these features with outcomes requires manual and heuristic aggregation across images. While promising, a significant hurdle in spatial pattern analysis is deciphering biologically and clinically relevant patterns from the complex architecture within tissue across various slides.

In recent literature, cell neighborhood (niche) analysis is emerging as a popular approach. This analysis pipeline typically consists of two primary steps by first identifying neighborhood features for each single cell using either a K-nearest-neighbor (KNN) graph or a defined radius, and then applying a clustering algorithm, such as k-means, Louvain, or Latent Dirichlet Allocation (LDA) [2, 6, 7, 13–15]. Seurat v5 [16] for instance, clusters cells using k-means based on similar cell type compositions, offering a straightforward niche analysis method. There are different variants of the approach depends on how to incorporate spatial information into the clustering process. UTAG [13] averages marker expression within the neighborhood for clustering, while BankSY [17] further refines this by combining local mean expression with individual cell expression. Spatial-LDA [14] incorporates spatial priors into clustering to allow proximity-closed cells to share similar cell neighborhoods. More recently, graph neural networks have been employed to discern cell neighborhood patterns, such as CytoCommunity [18]. However, deep learning methods like CytoCommunity require significant computational resources, posing challenges for individual labs, particularly for large-scale image analysis. Other studies adapt computational methods designed for spatial transcriptomics to analyze tissue imaging data [19–21], such as those intended for 10x Visium, face limitations due to high computational costs [17] and are generally restricted to single tissue sections with fewer spots [13, 19]. These methods struggle with large-scale images, like whole-slide multiplexed data containing millions of cells, and are challenging to adapt for modern imaging platforms like Nanostring CosMx and 10x Xenium.

Highly interpretable and scalable machine learning methods are in great need for analyzing molecular tissue imaging data. In this work, we propose *SpatialTopic*, a Bayesian topic model designed to identify and interpret spatial tissue architecture across various multiplexed images by considering both the cell types and their spatial arrangement(Figure 1A). We adapt an approach originally developed for image segmentation in computer vision [22], incorporating spatial information into the flexible design of regions (image partitions, analogous to documents in language modeling). Unlike standard image pixels, the basic units of analysis in multiplexed tissue images are cells, which are not uniformly distributed due to the complexity of human tissue samples, posing a unique challenge. To address these challenges, we refined the original model used for image segmentation by using a nearest-neighbor kernel function to boost computational efficiency, as well as a unique initialization strategy for robustness. In addition, we also provide an efficient C++ implementation of the spatial topic model in our R package *SpaTopic*.

**Fig. 1.**
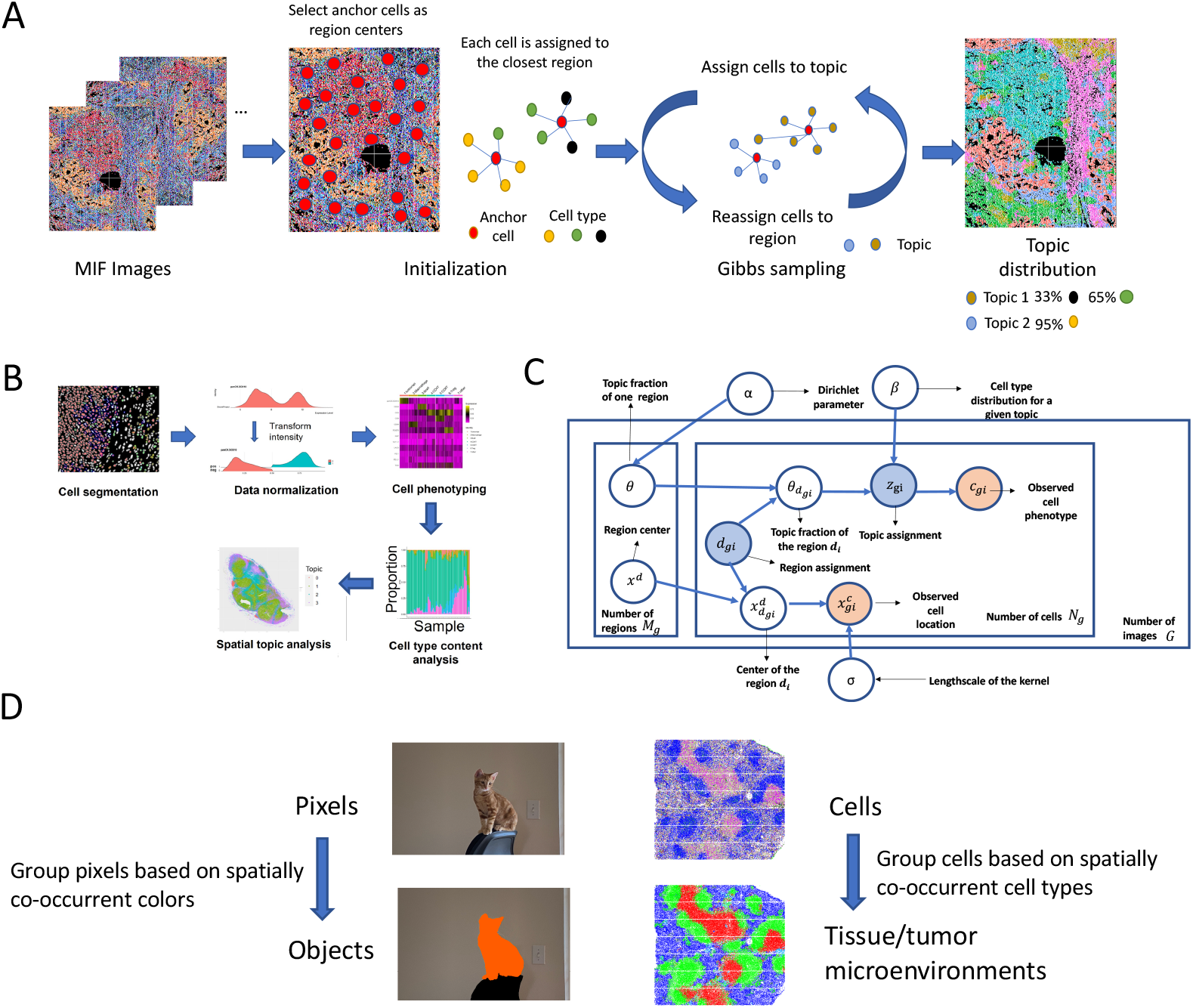
*Spatial Topic* unsupervisedly identifies distinct tissue microenvironments across images, utilizing topic model concepts in computer vision. A. Overview of *SpatialTopic. SpatialTopic* identifies biologically relevant topics across multiple images, while each topic is a distribution of cell types, reflecting the spatial tissue architecture across images. B. Image analysis pipeline designed for multiplexed immunofluorescence images. *SpatialTopic* is designed as a critical step for spatial analysis after cell phenotyping. C. Graphic representation for *SpatialTopic*. The observed and hidden variables are colored orange and blue accordingly. D. *SpatialTopic* groups cells in an unsupervised manner based on spatially co-occurrent cell types, similar to image segmentation based on spatially co-occurrent colors in the photo.

*SpatialTopic* offers a scalable solution for cell neighborhood and domain analysis on large-scale, multi-image datasets, efficiently handling data without requiring the extraction of cell neighborhood information for each individual cell – a process that becomes computationally demanding and inefficient with millions of cells. Unlike the rigid clustering strategies of other methods, *SpatialTopic* identifies ‘topics’—tissue microenvironment features—through a probabilistic distribution over cell types and *across* diverse tissue images using a generative model. We demonstrate our method can accurately identify and quantify interpretable and biologically meaningful topics from imaging data without human intervention. We also present multiple case studies encompassing tissue images from mouse spleen, non-small cell lung cancer, healthy lung, and melanoma tissue samples. Finally, we highlight an example of a TLS-like topic and its correlation with outcomes from *SpatialTopic* analysis across different platforms, as well as a multi-stage example showing dynamic changes in spatial tissue architecture across varying disease stages.

## Results

### Overview of *SpatialTopic*, a Bayesian probabilistic model for highly scalable and interpretable spatial topic analysis *across* multiplexed tissue images

*SpatialTopic* is designed as a flexible spatial analysis module within the current imaging analysis workflow (Figure 1B). Its main objective is to identify biologically meaningful topics *across* multiplexed images using unsupervised learning. Here, “topics” refer to latent spatial features defined by distinct cell type compositions within tissue microenvironment neighborhoods. *SpatialTopic* incorporates spatial data into a Latent Dirichlet Allocation model, assuming that each cell in an image arises from a mixture of spatially resolved topics, with each topic being a distribution over distinct cell types. Combining cell type information with spatial orientation, this method enables the automated and simultaneous detection of immunological patterns *across* multiple images. Subsequent analyses can further link these topics with patient data, such as treatment response and survival.

We adopt a Bayesian approach for inference to model the uncertainties inherent in tissue spatial patterns. *SpatialTopic* requires cell types and their locations as input, with the cell types determined by the users’ preferred phenotyping algorithm tailored to the specific marker panel of the dataset. The algorithm generates two key statistics for further analysis: 1) topic content, a spatially-resolved topic distribution over cell types, and 2) topic assignment for each cell within the images. After Gibbs sampling, the topic assignment of each cell is determined by the topic with the highest posterior probability. Cell types enriched in the same topic tend to be spatially correlated across images, leading to the identification of recurrent patterns of cell-cell interactions.

We developed an R package *SpaTopic* to efficiently implement the *SpatialTopic* algorithm as outlined in Figure 1A, which details the primary steps of the algorithm (See the Methods section). Figure 1C displays a graphical representation of the spatial topic model. The key inputs for *SpatialTopic* are the cell type annotations *C* and their locations *X* across all images. Here, *Z*_*gi*_ denotes the topic assignment, and *D*_*gi*_ indicates the region assignment of cell *i* in image *g*. Analogous to how computer vision algorithms segment images by spatially co-occurring pixel patterns with similar color, intensity or texture for object detection, *SpatialTopic* identifies topics as clusters of spatially co-occurring cell types (shown in Figure 1D), potentially corresponding to biologically meaningful cellular structures (e.g., tertiary lymphoid structure). The process involves the following steps:

- Initialization: Anchor cells are chosen as regional centers via spatially stratified sampling. For each image, a KNN graph is constructed between anchor cells and all other cells: For each cell, we retrieve its top *m* closest anchor cells. The initial region assignments of cells are made based on proximity to region centers.
- Collapsed Gibbs sampling: for every individual cell, there are two main steps per iteration:
  – Sample topic assignment *Z*_*gi*_ conditional on its region assignment *D*_*gi*_ and cell type *c*_*gi*_, as well as the topic distribution of the region *D*_*gi*_ and the cell type distribution of the topic *Z*_*gi*_.
  – Sample region assignment *D*_*gi*_ conditional on current topic assignment *Z*_*gi*_, distance of the cell 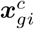 to the region center 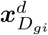, and the topic distribution of the region *D*_*gi*_. The spatial information is weakly incorporated with a kernel function.
- After Gibbs sampling, the output includes the posterior probabilities *Z*_*gi*_ of each cell and the per-topic cell type distribution 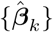. Each cell in the image is assigned to a topic with the highest posterior probability *P* (*Z*_*gi*_|*𝒞, 𝒳*).

We applied *SpatialTopic* to multiple datasets from diverse imaging platforms, including spatial proteomics data from Co-detection by Indexing (CODEX), Multiplexed ImmunoFluorescence (mIF), and Imaging Mass Cytometry (IMC) platforms, as well as spatial transcriptomics data from Nanostring CosMx (Table S1). In the next few sections, we apply *SpatialTopic* to analyze tissue imaging data from a variety of spatial molecular profiling platforms and benchmark analysis of *SpatialTopic* with other popular algorithms for spatial domain/niche analysis, including Seurat v5 [16], Spatial-LDA [14], CytoCommunity [18], UTAG [13], and BankSY [17] (Table S2). The benchmark datasets contain between 0.1 to 1 million cells per image; making it challenging to apply methods with high computational costs. In contrast, *SpatialTopic* processes these large-scale images within just a few minutes.

### *SpatialTopic* identifies global and local spatial features of human lung cancer tissue with higher precision and interpretability

We applied our method to a single non-small cell lung cancer (NSCLC) tissue image generated using a 960-plex CosMx RNA panel on the Nanostring CosMx Spatial Molecular Imager platform, which is publicly available on the Nanostring website. We selected a Lung5-1 sample containing approximately 100,000 cells, with 38 cell types annotated using Azimuth [23] based on the human lung reference v1.0 (Figure 2A).

**Fig. 2.**
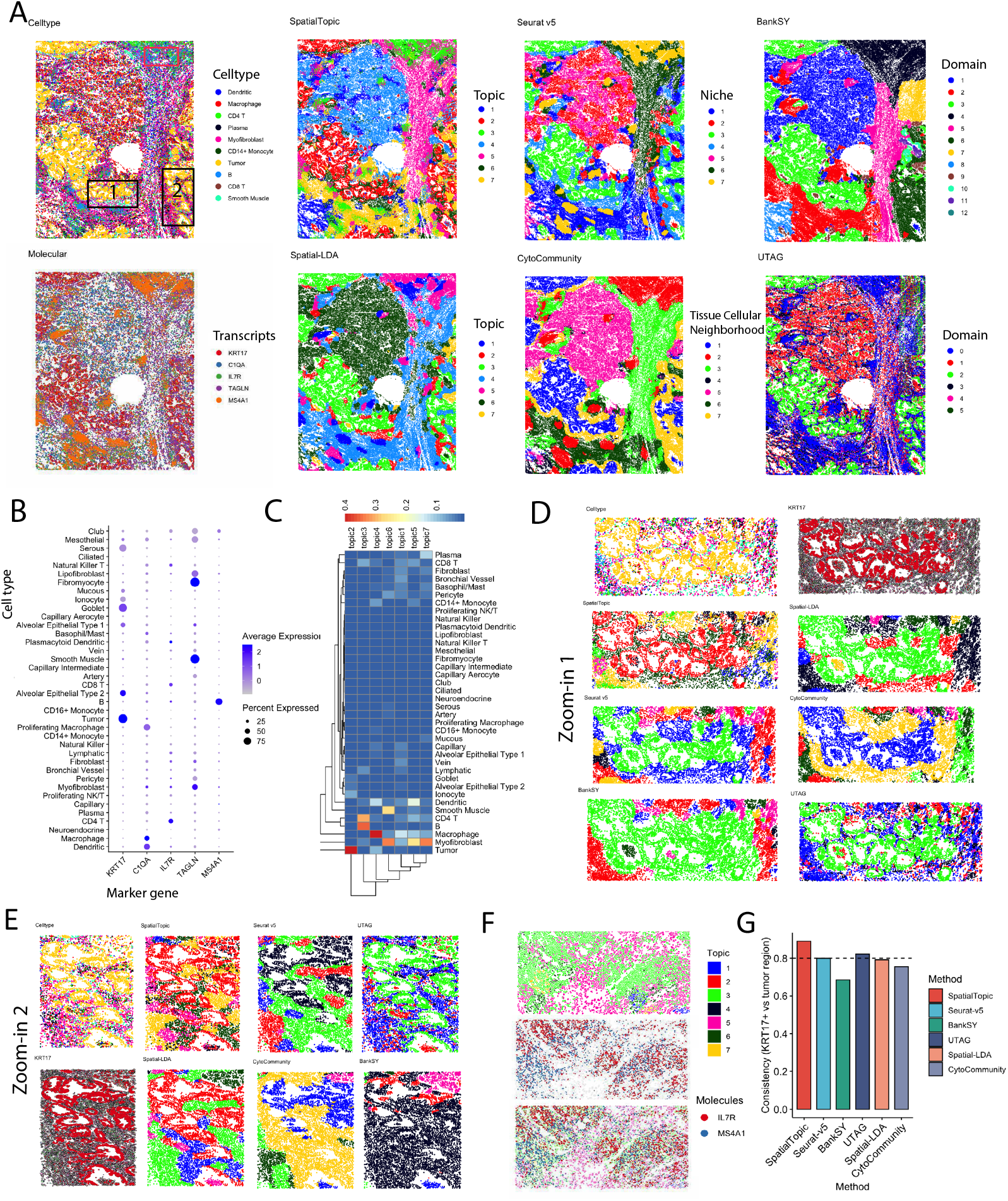
*SpatialTopic* better detects tumor microenvironment in a nanostring human non-small cell lung cancer tissue image. A. We compare *SpatialTopic*, Seurat v5, Spatial-LDA, CytoCommunity, BankSY, and UTAG results on the human lung tumor tissue samples. We also visualize the distribution of the top 10 most abundant cell types and five unique mRNA molecules (*KRT17, C1QA, IL7R, TAGLN, MS4A1*), showing the tissue architecture. We only show up to a total of 20k molecules due to limitations in visualization. B. Dot plots showing gene marker expression across all 38 annotated cell types. *KRT17, C1QA, IL7R, TAGLN*, and *MS4A1* are marker genes for tumor, macrophage, CD4 T, stroma, and B cells, respectively. C. Heatmap shows per-topic cell type composition. Topic 2 represents tumor regions. The other topics represent distinct immune-enriched stroma regions, including topic 3, which captures the lymphoid structure in the lung tissue consisting of B cells and CD4 T cells, and topic 4, which is a macrophage-enriched stroma region. D. and E. *SpatialTopic* can better capture the local structure of the lung tumor tissue. F. Topic 3 (green) captures the lymphoid structures, consistent with the distribution of *IL7R* (CD4 T cells, red) and *MS4A1* (B cells, blue). G. We compare the consistency of different results, presenting the percentage of cells in the identified tumor domains expressing the *KRT17* gene. *SpatialTopic* and UTAG generally show higher consistency than other methods.

To illustrate the general tissue architecture, Figure 2A displays the distribution of the top 10 main cell types and the expression patterns of key genes including *KRT17, C1QA, IL7R, TAGLN, MS4A1*. These genes serve as markers for tumor cells (*KRT17*), macrophages, CD4 T cells, stroma cells, and B cells, respectively (Figure 2B). Our results demonstrate that *SpatialTopic* identified seven distinct topics from the complex image (Figure 2A), with each topic representing a unique spatial niche characterized by a specific cell-type composition, as detailed in Figure 2C. For example, Topic 2 is predominantly composed of tumor cells, indicating the tumor region in the image, while other topics correspond to distinct immune-enriched stromal regions. Topic 4 represents a stromal region enriched with macrophages. Notably, Topic 3 captures tertiary lymphoid-like structures in the lung tissue, consisting of B cells, CD4 T cells, and smaller proportions of dendritic cells and CD8 T cells. This composition aligns with the current understanding of cell types in tertiary lymphoid structures, which are strong predictive biomarkers associated with a good prognosis and response to immunotherapy in non-small cell lung cancer [24].

We compared results from *SpatialTopic* with Seurat v5, Spatial-LDA, CytoCommunity, BankSY, and UTAG. BankSY and UTAG directly use cell-level gene expression as input, whereas the other four methods, including *SpatialTopic*, rely on cell-type annotations. All methods can detect the global structure of the image and classify tumor and stromal regions. However, BankSY and UTAG appear to miss the lymphoid structure, likely because they do not use the detailed information provided by cell-type annotations. Reference-based cell type annotation typically offers more detailed information and can be more robust for noisy data when matched with a single-cell reference [25, 26]. SpatialTopic distinctly identified the lymphoid structure as Topic 3, comprising a mix of CD4 T cells and B cells (Figure 2F). Additionally, when we focused on two local tumor tissue regions (Figures 2D and 2E), *SpatialTopic* identified the tumor region with higher precision (Topic 2), more consistently matching the expression pattern of *KRT17*, a lung cancer marker gene. *SpatialTopic* and UTAG are the only two methods showing the consistency (between tumor domain and *KRT17* expression) higher than 0.8 across the entire image (Figure 2G), which aligns with the visual measure in Figure 2D and 2E.

### *SpatialTopic* identifies tertiary lymphoid structures from whole-slide melanoma tissue imaging

We applied *SpatialTopic* to a whole-slide melanoma tissue image obtained from our internal multiplexed immunofluorescent (mIF) imaging platform, which uses a 12-plex marker panel [27]. This analysis covered a whole-slide soft tissue image containing 0.4 million cells, annotated into seven major cell types (CD4 T cells, Tumor/Epithelial, B cells, CD8 T cells, Macrophages, Regulatory T (Treg) cells, and Others). The categorization was based on the expression of six lineage markers: CK/SOX10, CD3, CD8, CD20, CD68, and Foxp3. Cells were annotated as ‘Other’ if they showed negative expression for all six markers.

Despite using fewer markers compared to the Nanostring CosMx platform, *SpatialTopic* identified five distinct topics (Figure 3A): Topic 1 (tumor), Topic 2 (CD4 immune zone), Topic 3 (stroma), Topic 4 (immune-enriched tumor-stroma boundary), and Topic 5 (tertiary lymphoid structures). The tissue structures revealed by these topics visually correspond to the histological pattern seen in the coregistered H&E image (Figure 3B) and the merged raw mIF images (Figure 3C) with three key markers: CD3 (T cells), CD20 (B cells), and PANCK/SOX10 (tumor cells). Figure 3D demonstrates that topic 5 (tertiary lymphoid structures) mainly consists of B cells, CD4 T cells, a few CD8 T cells, and Treg cells, consistent with the TLS-like pattern identified in the Nanostring dataset discussed earlier. Due to the lack of a dendritic cell marker in the mIF dataset, dendritic cells could not be identified and included in topic 5. This analysis demonstrates that *SpatialTopic* can consistently detect the same biologically relevant patterns across various tumor tissues and imaging platforms, which may be clinically significant, as tertiary lymphoid structure have been recognized as a promising biomarker for cancer immunotherapy.

**Fig. 3.**
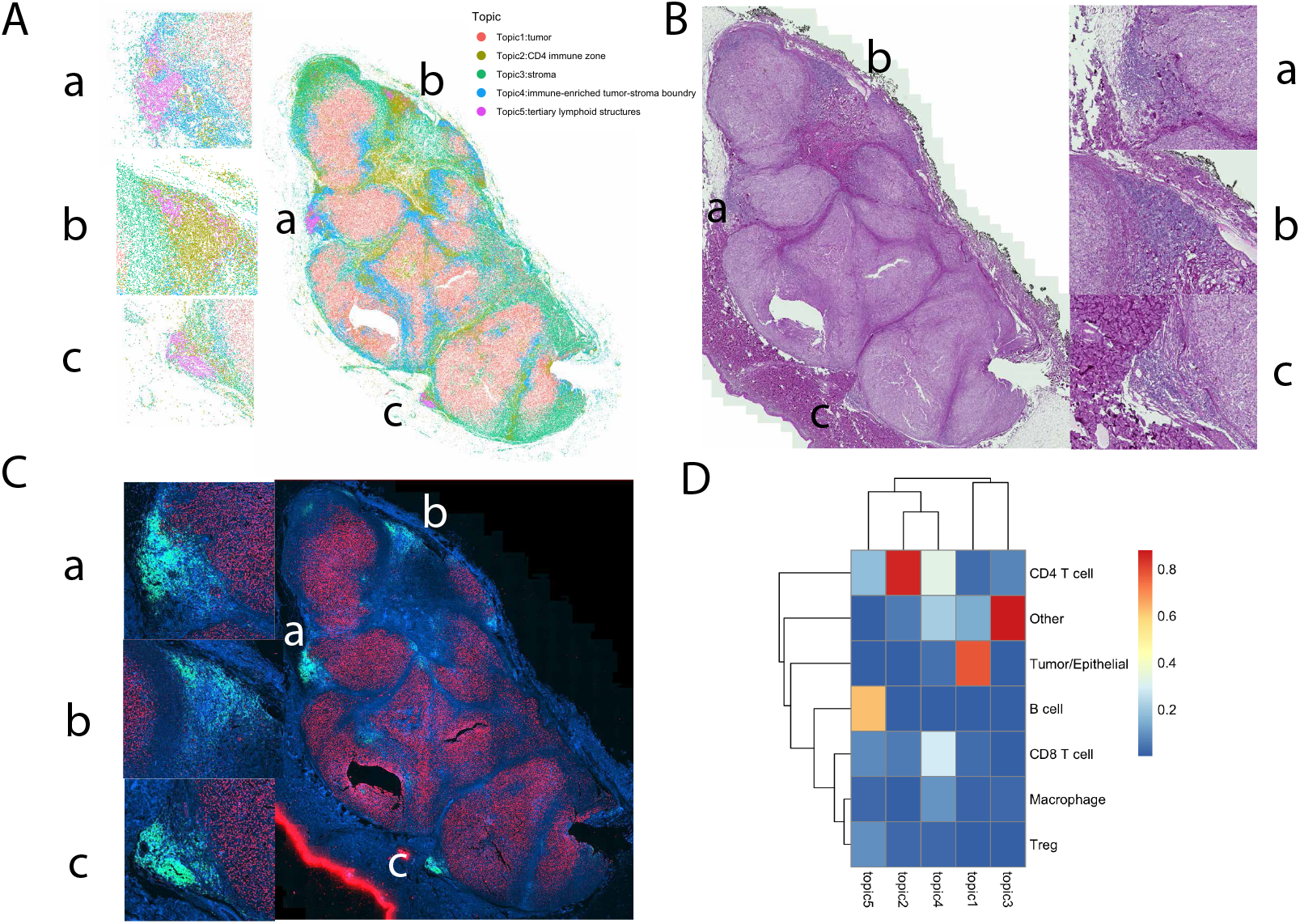
*SpatialTopic* identifies tertiary lymphoid structures from a whole-slide melanoma tissue sample. A. *SpatialTopic* identifies five topics from the whole-slide melanoma tissue sample: topic 1 for tumor region, topic 2 for CD4 T cell region, topic 3 for stroma region, topic 4 for immune-enriched stroma-tumor boundary, topic 5 for the potential tertiary lymphoid structures, with three Region of Interests (ROIs) highlighted in the subfigures. B. H&E staining images for the whole-slide melanoma tissue sample and three ROIs. C. Merged mIF image for the whole-slide melanoma tissue sample and three ROIs with three channels: PANCK/SOX10 (red), CD3 (royal blue), and CD20 (green). D. Heatmap shows per-topic cell type composition for the five topics identified by *SpatialTopic*. Topic 5 (tertiary lymphoid structures) mainly consists of B cells and CD4 T cells, with a small proportion of CD8 T cells and regulatory T (Treg) cells.

### *SpatialTopic* recovers spatial domain from cell type spatial organization in healthy lung tissue

We further demonstrate that *SpatialTopic* can effectively distill signals from noisy cell type annotations and identify clear tissue architecture based solely on the spatial arrangement of cells. To illustrate this, we applied *SpatialTopic* to the IMC dataset from the UTAG paper [13], which includes 26 small regions of interest (ROIs) images from healthy lung tissue. For comparison, we used the UTAG result provided in the paper [13] without rerunning UTAG.

Our analysis shows that *SpatialTopic* can recover tissue architectures directly from the spatial distribution of cell type annotations, yielding results consistent with manual annotations (Figure 4A). *SpatialTopic* performs comparably to UTAG using only cell type annotations (Figure 4B), as indicated by the adjusted Rand index, which shows similar performance levels. Additionally, Figure 4C illustrates the topic content and cell type composition for each topic identified by *SpatialTopic*. This demonstrates *SpatialTopic*’s capability to perform domain analysis without discarding existing cell type annotations, offering valuable flexibility for datasets with cell-type annotations or for incorporating any existing cell-type annotation method. Unlike UTAG, which learns spatial tissue architecture directly from cell features due to noisy cell type annotations, we demonstrate that *SpatialTopic* can effectively identify tissue architecture from these annotations. Thus *SpatialTopic* is a robust alternative that leverages existing data without the need for additional cell-level features.

**Fig. 4.**
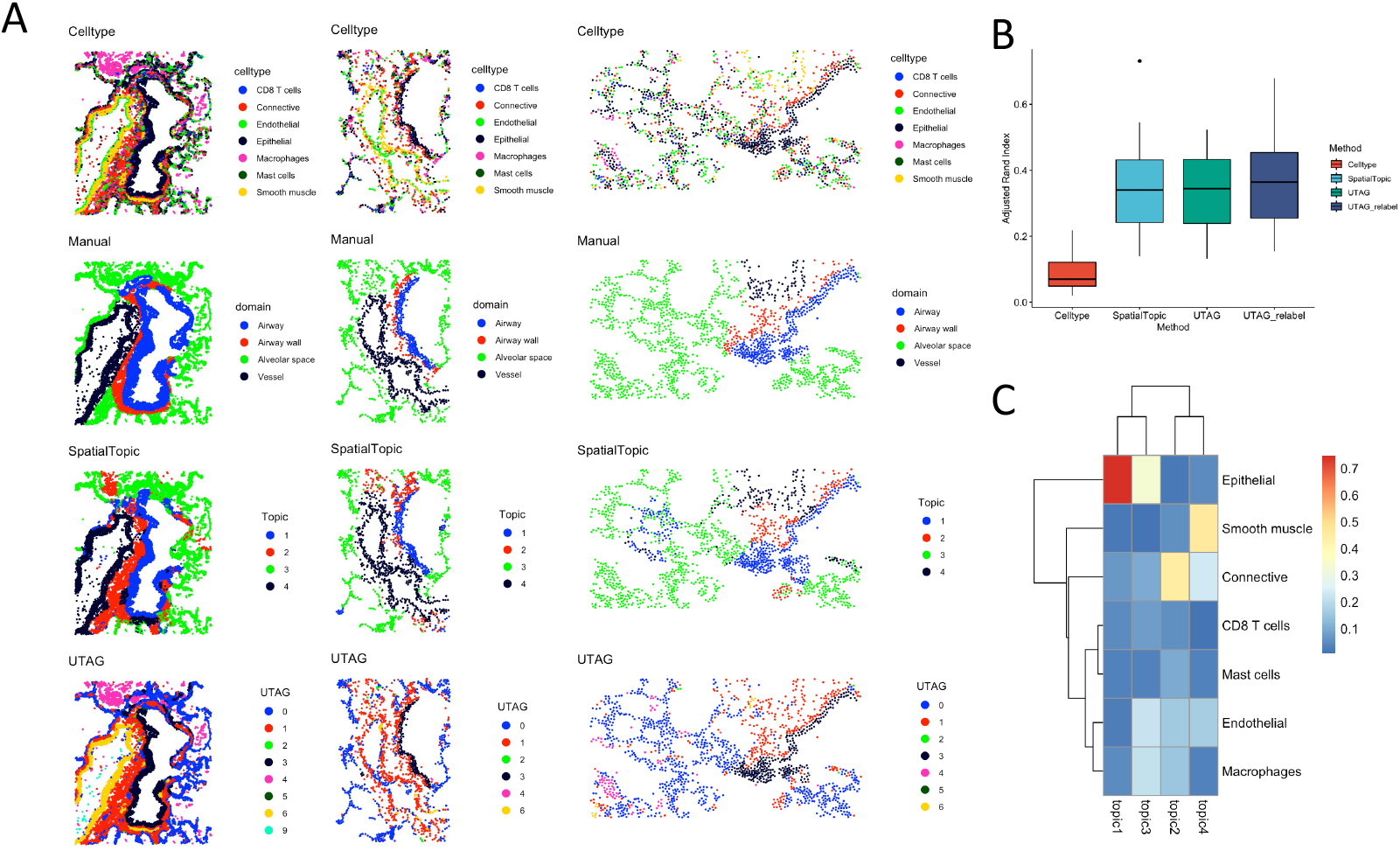
*SpatialTopic* recovers spatial domain architecture from cell type spatial layout in healthy lung. A. Spatial distribution of cell type annotations, manual domain annotations, and spatial domains recovered by *SpatialTopic* and UTAG from the healthy lung tissue samples. B. Consistency comparing manual spatial domain to the cell type annotation, *SpatialTopic*, and UTAG with and without ad hoc relabel across 26 images. UTAG results are obtained from the original publication. C. Heatmap shows per-topic cell type composition for the four topics identified by *SpatialTopic*.

### *SpatialTopic* identifies disease-specific topics and tracks topic evolution in mouse spleen over disease progression

We also applied *SpatialTopic* to a CODEX mouse spleen dataset [2] to demonstrate its proficiency in identifying spatial topics across multiple images. This dataset includes nine images: three control normal BALBc spleens (BALBc 1-3) and six MRL spleens (samples 4-9) at varying disease stages—early (MRL 4-6), intermediate (MRL 7-8), and late (MRL 9) (Figure 5A). Using a 30-plex protein marker panel, the study identified 27 major splenic-resident cell types across the nine tissue images. We use the cell type annotation in the original paper [2].

**Fig. 5.**
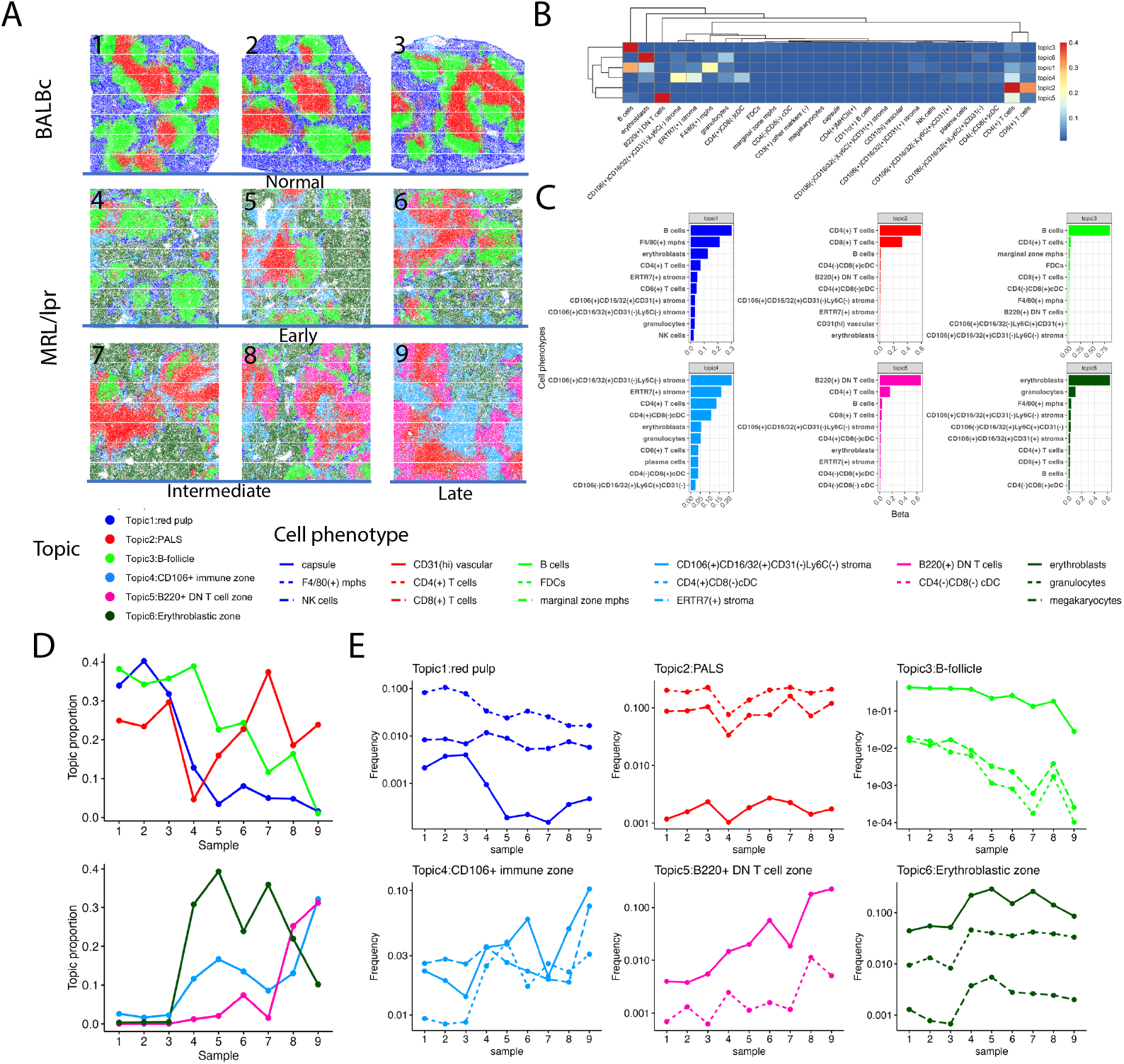
*SpatialTopic* captures main dynamics in tissue architecture of normal and diseased mouse spleen. A. Six topics were identified by *SpatialTopic* across nine mouse spleen samples representing normal (BALBc 1-3) and different disease stages: early (MRL 4-6), intermediate (MRL 7-8), and late (MRL 9). B. Heatmap shows per-topic cell type composition for the six main topics identified by *SpatialTopic*. Based on cell type compositions, the first three topics are labeled as red pulp, PALS (periarteriolar lymphoid sheath), and B-follicle in normal mouse spleen tissue, while the other three topics are unique to the disease stages. C. Barplots shows the top 10 per-topic cell types for the six main topics identified by *SpatialTopic*. D. Dynamic change in the topic proportion of the six topics during disease progression. Normal spleen samples are primarily characterized by topics 1, 2, and 3, which reflect red pulp (mixed of B cells, erythroblasts, and F4/80(+) mphs), PALS (mixed of CD8 T cells and CD4 T cells), and B-follicle (most B cells) respectively. There is an increase in Topic 6 and depletion of Topic 1 in MRL samples, representing much fewer B cells and F4/80(+) mphs but more granulocytes and erythroblasts in the red pulp regions. Topic 5 (mainly B220(+) DN T cells and CD4(+) T cells) is enriched in tissue at late disease stage. E. Dynamic change of key immunological cell types within each topic, identified by FREX(omega = 0.9) and lift metrics (See Figure S5).

*SpatialTopic* identified six topics from approximately 0.7 million cells across the nine images, highlighting the dramatic changes in spatial tissue structures associated with disease progression from normal spleen to spleen tissue at different disease stages (Figure 5A). Figure 5B and 5C highlight per-topic cell type compositions, aiding in labeling each topic. The normal spleen tissue samples predominantly comprised three topics: Topic 1 (red pulp), Topic 2 (periarteriolar lymphoid sheath, PALS), and Topic 3 (B-follicle). Figures S2 and S3 show the cell type distribution and domain annotations from the original paper, demonstrating *SpatialTopic*’s ability to capture the main structures consistent with these annotations, as compared to other methods (Figures S2 and S4). With an increasing number of topics, *SpatialTopic* also successfully delineated the marginal zone from the B-follicle (Figure S2).

Topics identified by *SpatialTopic* were comparable across normal and diseased spleens, allowing us to identify condition-specific topics and quantify changes in topic proportions as the disease progressed. In contrast to normal spleens, MRL spleens showed a decrease in B cells and F4/80(+) macrophages but an increase in granulocytes and erythroblasts within the red pulp region, indicating inflammation or systemic infection in the spleen tissue. This shift was marked by the predominance of Topic 6 in MRL spleens, superseding Topic 1 (red pulp). Topic 4 emerged in the mouse spleen tissue affected by autoimmune disease, characterized by a high abundance of CD106+ stroma cells, an indicator for leukocyte recruitment to inflamed areas. Thus, this topic also shows a high concentration of immune cells, including CD4 and CD8 T cells. Unique to MRL spleens, Topic 5 is characterized by an enrichment of B220+ double negative (DN) T cells and conventional CD4 T cells, predominating in tissues during advanced stages of the disease, indicating a shift in the immune cell landscape. These dynamics indicate immune surveillance or dysregulation in the spleen tissue as disease progression [2].

Furthermore, *SpatialTopic*’s capability to identify topics based on the spatial proximity of cell types suggests that cell types grouped within the same topic are likely close to each other and prone to interaction. Figure 5D illustrates the changes in topic proportions throughout the course of the disease progression. The distinct contributions of cell types to each topic are highlighted in Figure 5E, selected based on their specific composition and evaluated based on the lift and FREX metrics [28, 29] (Figures S5). Cell types are clustered into topics that exhibit similar dynamics across different slides.

### *SpatialTopic* is highly scalable on large-scale modern images

To benchmark the scalability of *SpatialTopic* as the number of cells in images increases, we conducted tests using simulated datasets of varying scales. Figure 6A shows that *SpatialTopic* significantly outperforms Seurat v5 in terms of scalability with an increasing number of cells within a single image. Moreover, as demonstrated in Figure 6B, *SpatialTopic* shows high efficiency on real large-scale imaging datasets, requiring less user time compared to other methods. For example, on the Nanostring CosMx NSCLC image with approximately 0.1 million cells, *SpatialTopic* runs within one minute on a standard MacBook Air, a performance unbeatable by other methods.

**Fig. 6.**
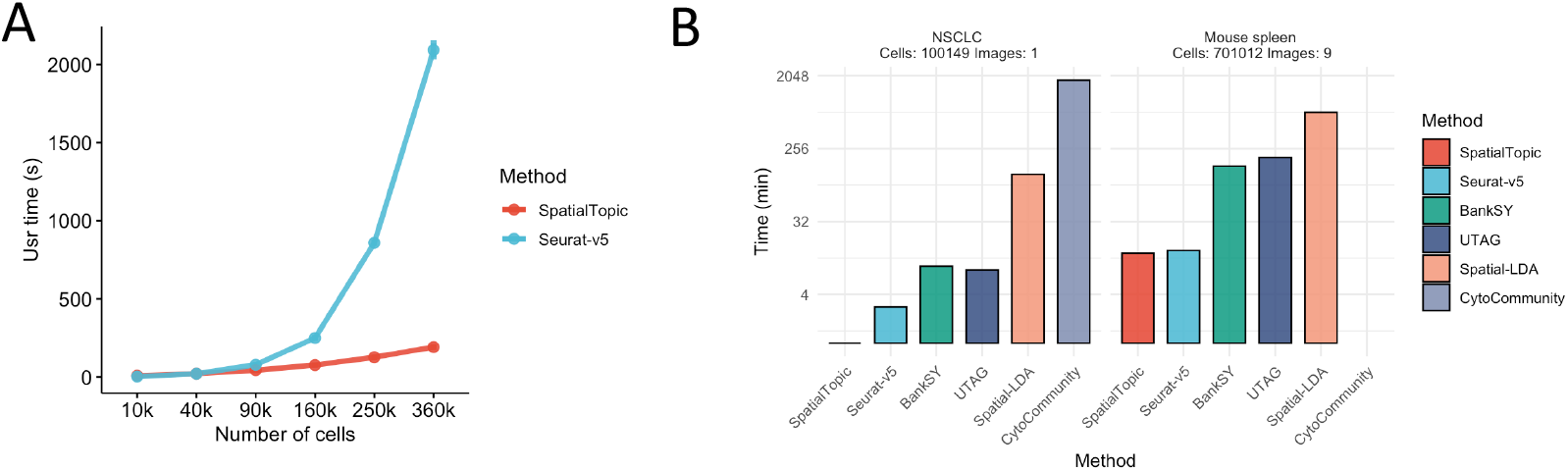
*SpatialTopic* is scalable to large-scale images and can be run on a regular laptop within minutes. A. Runtime of SpatialTopic (region radius r = 60) and Seurat-v5 on simulated datasets with increasing cell numbers on a single image. B. Runtime of *SpatialTopic*, Seurat-v5, BankSY, UTAG, Spatial-LDA, and CytoCommunity on large-scale NSCLC and mouse spleen datasets. All methods were benchmarked on a standard MacBook Air (M2, 2022) unless exceeding the memory limitation.

Across all datasets, *SpatialTopic* ranks in the highest tier with Seurat v5, while BankSY and UTAG fall into a second tier due to their reliance on similar, but less optimized, strategies. CytoCommunity, limited by its dependency on GPU support and memory demands, was run with reduced epochs and CPU-only for the NSCLC dataset, which compromised its performance and underscored its impracticality for labs without extensive computing resources on large-scale imaging analysis. Additionally, on images with cells more than 0.1 million, both UTAG and CytoCommunity required running on high-performance computing server, due to their high memory demands. In contrast, *SpatialTopic* is highly scalable on large-scale imaging analysis, with all analysis done within minutes on a standard laptop.

## Discussion

In summary, we introduced *SpatialTopic*, a spatial topic model designed to identify and quantify biologically relevant topics across multiple multiplexed tissue images. This represents a novel approach to applying language modeling techniques to decipher the tissue microenvironment from tissue imaging data. *SpatialTopic* stands out as one of the few unsupervised learning methods capable of discerning clinically relevant spatial patterns [13, 17, 19]. Unlike other methods that rely on hard clustering strategies for analyzing samples, *SpatialTopic* is a probabilistic model-based approach using Bayesian inference methods to identify complex tissue architectures. The model generates two key outputs: The first of these, the *topic content* maps the cell type composition in spatial niches, allowing direct interpretation of the corresponding topic (e.g., TLS); The second output, *topic assignment* for each single cell allows the quantification of each topic in individual tissue samples for subsequent association analysis with patient outcome. Application to multiple datasets along with benchmark analysis show that *SpatialTopic* achieves higher precision in defining global and local spatial niches and higher sensitivity at capturing complex structures such as TLS. Notably, our method is highly scalable to large-scale imaging data with efficient runtime, handling millions of cells on a standard laptop.

*SpatialTopic* is designed as a flexible spatial analysis module within the current imaging analysis workflow. A standard image analysis pipeline includes cell segmentation, data normalization/batch correction, cell phenotyping/clustering, and the analysis of cell type content and spatial relationships. Downstream statistical analysis typically starts with cell-level metadata derived from image analysis. Due to varied marker panels and molecular imaging platforms, a one-size-fits-all solution for cell phenotyping across diverse platforms seems unlikely. In practice, we find that reference-based cell annotation works best on single-cell imaging data, rather than unsupervised clustering (data not shown). *SpatialTopic* does not specify any upstream method, and thus can be seamlessly integrated with other cell phenotyping modules tailored for datasets from different platforms. This design offers users adaptability, accommodating datasets from different panel designs.

In our proposed analysis pipeline for imaging data, we separate cell phenotyping from cell neighborhood/domain analysis for image-based spatial data, with *SpatialTopic* directly taking cell types as input. This key difference sets *SpatialTopic* apart from UTAG and BankSY, which use protein/gene expression as input for niche/domain analysis. UTAG performs dimension reduction before message passing, while BankSY engineers new spatial features for each cell before dimension reduction. We propose that treating cell phenotyping and neighborhood/domain analysis as distinct steps is a better analysis strategy for datasets generated by image-based technology with selected marker panels. Using cell type annotations as input for cell neighborhood analysis enhances the interpretability of different tissue microenvironments and undoubtedly increases the computational efficiency when analyzing largescale images. The performance of *SpatialTopic* may rely on the accuracy of cell phenotyping. A better strategy for cell phenotyping is to annotate cells directly from cell images instead of using summary statistics, such as mean marker expression or gene count data. As part of the analysis pipeline, we are developing an image-based deep learning method for cell phenotyping, incorporating subcellular information, as well as domain knowledge [30].

For multi-sample analysis, addressing the batch effect is a key challenge. Our proposed analysis pipeline seeks to mitigate the batch effect during cell phenotyping using a reference-based cell phenotyping method. For spatial transcriptomics data, a supervised classification method with a reliable single-cell reference can mitigate batch effects and inherent noises in the imaging data. Batch effect is more critical for algorithms that directly consider the gene expression data as input. When analyzing the mouse spleen dataset, we used Combat [31] for batch correction across multiple images before applying UTAG and BankSY. However, Combat appears to over-correct for batch effects (Figure S4), thus failing to distinguish between normal and diseased red pulp tissue. This might stem from the substantial differences between normal and diseased tissues.

Modern datasets from platforms like 10x Xenium and Nanostring CosMx require scalable computational methods to handle their size and complexity. Existing spatial domain analysis methods, originally designed for 10x Visium spatial transcriptomics data and optimized for datasets with thousands of cells or spots per slide, find it challenging to handle these more advanced, datasets with millions of cells per image. *SpatialTopic* meets this need by efficiently managing neighborhood calculations and constructing the KNN graph only among *m* anchor cells instead of all *n* cells in the image. This reduces the time complexity from *O*(*n* log *n*) to *O*(*m* log *m*), where *m* ≪ *n*. Additionally, *SpatialTopic* maintains linear time complexity relative to the number of cells and iterations with collapsed Gibbs sampling and uses an approximate fast approach for constructing the KNN graph. These optimizations ensure *SpatialTopic*’s computational efficiency, making it accessible on standard laptops and practical for analyzing large-scale imaging data from platforms like the 10x Xenium and Nanostring CosMx.

Moreover, advances in technology now enable the quantification of immune cell spatial diversity and the characterizing of tumor microenvironments in 3D tissues [32]. While *SpatialTopic* can be adapted to infer immunological topics from 3D tissue, a refined strategy is needed to select anchor cells in the 3D spaces, as the spatial information obtained by *SpatialTopic* primarily stems from the relationships between the anchor cells and other cells. Incorporating a hierarchical Dirichlet prior on topic distributions across regions would allow regions within the same image to share priors while differing across images. Furthermore, optimizing the initialization strategy is needed when applying *SpatialTopic* to extremely large datasets with hundreds of images. These improvements would broaden the applicability of *SpatialTopic*.

## Methods

### SpatialTopic

#### Notations

We assume there are total *V* cell types that contribute to *K* different tissue microenvironments (topics) across *G* multiplexed images. Let *c*_*gi*_ be the *i*th cell at the location 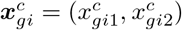, *g* = 1, 2, …, *G, i* = 1, 2, …, *n*_*g*_, on the *g*th image with total *n*_*g*_ cells. Let *c*_*gi*_ = *v* if the cell has been classified to the *v*th cell type. Let 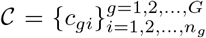 and 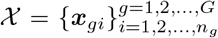 denote all observed cell types and cell locations across all *G* images.

#### Model

In a conventional LDA model, each image is treated as an individual document, employing a bag-of-words approach without accounting for spatial information. This approach is similar to our prior work on longitudinal flow cytometry data analysis [28]. Here, in order to incorporate spatial information within images, we introduce a spatial topic model, *SpatialTopic*, integrating spatial data into the foundational LDA framework. This spatial topic framework was first proposed for image segmentation [22], instead of viewing each image as a singular document, we treat each image consisting of densely placed overlapping regions (documents). Unlike the conventional LDA model where relationships between documents and words are known and fixed, the word-document relationship here is unknown: each cell (word) is flexible to be assigned to all possible regions (documents). This flexible region (document) design allows us to identify spatial structure with irregular shape.

For *SpatialTopic*, we introduce a new hidden variable *D*_*gi*_ to denote cell region (document) assignment. Thus, each cell is associated with two hidden variables: the latent topic assignment *Z*_*gi*_ *∈ {*1, 2, …, *K}* and the latent region assignment *D*_*gi*_ *∈ {*1, 2, …, *M}, M* = Σ_*g*_ *M*_*g*_, where *M*_*g*_ denote the number of regions on the image *g*. During the initialization, we pre-selected anchor cells as region centers. Let 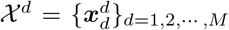 be the set of all M region centers across all images. Let ***θ***_*d*_ be the proportion of region *d* over *K* topics and ***β***_*k*_ be the proportion of topic *k* over *V* cell types. Hyperparameters ***ψ*** and *α* specify the nature of the Dirichlet priors of *{****β***_*k*_ *}* and *{****θ***_*d*_*}*, respectively.

Then we are ready to describe our generative model:

- For each topic k, sample ***β***_*k*_ (topic weights over *V* cell types) from a Dirichlet prior ***β***_*k*_ ∼ *Dir*(***ψ***).
- For each image region *d* (centered at 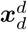), sample topic proportion ***θ***_*d*_ ∼ *Dir*(***α***)
- For each cell, the *i*th cell in the image *g*:
  – Sample its region assignment *D*_*gi*_ from a uniform prior over possible documents (regions) in the image *g*.
  – Sample the location 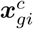 conditional on its region assignment *D*_*gi*_ with a kernel function based on the distance between the cell location 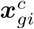 and the region center 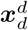.

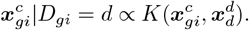
  – Sample topic assignment *Z*_*gi*_|*D*_*gi*_ = *d ∼ Multi*(***θ***_*d*_, 1).
  – Sample cell type *c*_*gi*_|*Z*_*gi*_ = *k ∼ Multi*(***β***_*k*_, 1).

Hyperparameters ***α*** and ***ψ*** should be chosen based on the belief on *{****θ***_*d*_*}* and *{****β***_*k*_ *}* in a Bayesian perspective. In our application, both ***α*** and ***ψ*** are set very small by default (default: *α*_*k*_ = .01, ∀*k*; *ψ*_*v*_ = .05, ∀*v*) to encourage the sparsity in region-topic distributions *{****θ***_*d*_*}* and topic-celltype distributions *{****β***_*k*_ *}*.

#### Nearest-neighbor Exponential Kernel

The flexible relationships between regions and cells in *SpatialTopic* allow each cell to be assigned to any one of its proximate regions. We employ a nearest-neighbor Gaussian kernel to capture the spatial correlation between cells and their respective regions, as previously used in the nearest-neighbor Gaussian process [33]. For computational efficiency, especially with large-scale images, we restrict our consideration to the top nearest-neighbor regions for each cell. Let 𝒩 (***x***_*gi*_) ⊂ *𝒳*_*d*_ be the collection of *m* closed region centers to the cell ***x***_*gi*_ (default: *m* = 5). In practice, the commonly used squared exponential Gaussian kernel function decays too rapidly. This rapid decay often results in cells predominantly being linked to their closest region, irrespective of their cell types. Let *σ* be the lengthscale that controls how fast correlation decays with distance in the kernel function. Thus, drawing inspiration from [34], instead of the squared exponential kernel, we used the following exponential kernel,

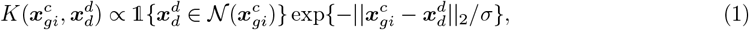

where 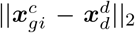 represents the Euclidean distance between the cell location 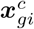 and the region center 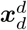. We fix *σ* for computational efficiency, but it can also be sampled during the Gibbs sampling. Increasing *σ* would reduce the strength of the spatial correlation, resulting in a diminished spatial effect when assigning cells to regions.

#### Collapsed Gibbs Sampling

We use collapsed Gibbs Sampling for model inference. The collapsed Gibbs Sampling algorithm was first introduced as the Bayesian approach of Latent Dirichlet Allocation [35]. This method’s comprehensive derivation and implementation can be found in the paper [36]. Similar to [22], we further adapted and extended the algorithm for our proposed spatial topic model. It’s noteworthy that during the collapsed Gibbs sampling process, the parameters ***β***_*k*_ and ***θ***_*d*_ are integrated out and are not explicitly sampled. Instead, our focus is on the two hidden variables associated with each cell: the topic assignment *Z*_*gi*_ and the region (or document) assignment *D*_*gi*_. These variables undergo iterative sampling using the collapsed Gibbs Sampler:

1. Sample topic assignment *Z*_*gi*_ conditional on region assignment *D*_*gi*_ with [35]

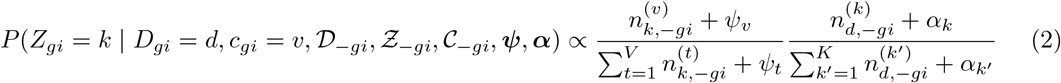

where 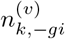 refers the number of times that cell type *v* has been observed with topic *k* and 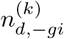 refers the number of times that topic *k* has been observed in region *d*, both excluding the current cell

*gi*, the *i*th cell on the *g*th image. The first ratio expresses the probability of cell type *v* under topic *k*, and the second ratio expresses the probability of topic *k* in region *d*. 𝒟_−*gi*_, 𝒵_−*gi*_, and 𝒞_−*gi*_ denote collections of 𝒟, 𝒵, and 𝒞 excluding cell *c*_*gi*_.

2. Sample *D*_*gi*_ conditional on *Z*_*gi*_ with

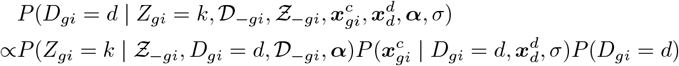

According to [36], *P* (*Z*_*gi*_ = *k* | *𝒵*_−*gi*_, *D*_*gi*_ = *d, 𝒟*_−*gi*_, ***α***) can be obtained by integrating out ***θ***_*d*_, that

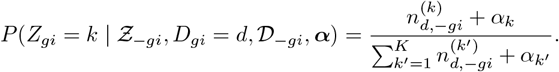

We can further omit *P* (*D*_*gi*_ = *d*) due to uniform prior. Thus *D*_*gi*_ can be sampled based on the following conditional distribution:

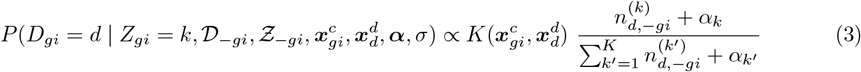

##### Algorithm 1

Collapsed Gibbs Sampling for *SpatialTopic*

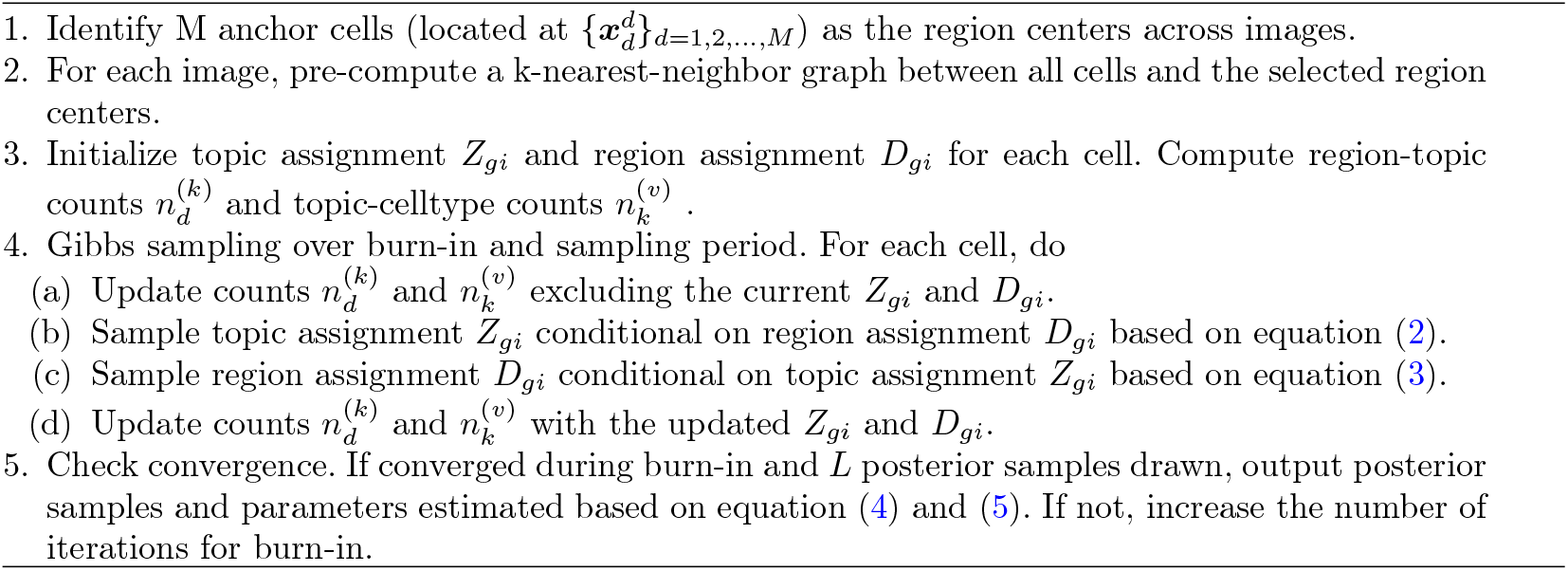

#### Initialization

During the initialization, we employ a spatially stratified sampling approach to randomly select anchor cells from each image, which will serve as region centers. The number of anchor cells selected from each image is determined by a predetermined region radius *r* (default: *r* = 400), as well as the image size. The radius should be set with the consideration of the image resolution and complexity of the images, and an adequate number of cells are expected within each region since it is crucial for estimating topic distribution ***θ***_*d*_ precisely. In practice, for whole-slide imaging, we expect at least 100 cells per region on average. For each individual image, an *m*-nearest-neighbor graph will be constructed between all cells and the chosen anchor cells. For computational efficiency, distances between each cell and its top *m*-nearest anchor cells will be pre-computed before Gibbs sampling.

The performance of *SpatialTopic* depends on anchor cells selected in the initialization, especially on images with highly complex spatial structures. Thus, we take a warm start approach rather than starting Gibbs sampling from a random initialization. This involves running multiple Gibbs sampling initializations (default: ninit = 10), each having a unique set of anchor cells. After a few iterations (default: niter init = 100), only the one with the highest log-likelihood is retained and continued.

#### Implementation

We implemented *SpatialTopic* in Rcpp and made it an R package *SpaTopic* (officially available on CRAN after Jan 17, 2024). The complete algorithm is shown in Algorithm 1. For the Gibbs sampling, we have set the default parameters as follows: iter = 200, burnin = 1000, thin = 20 (200 Gibbs sampling draws are made with the first 1000 iterations discarded and then every 20th iteration kept). We can infer topic distributions across all images using the posterior samples drawn from the Gibbs sampling. For each of these posterior samples, the predictive distributions of parameters *{****β***_*k*_ *}* and *{****θ***_*d*_*}* are obtained as follows:

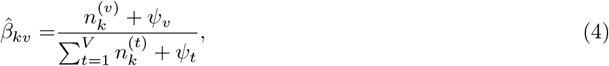

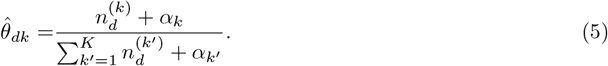

Moreover, we also keep the posterior distribution of *Z*_*gi*_ from all posterior samples for each individual cell. Notably, *D*_*gi*_ has been marginalized during this process and each cell in the end is assigned to the topic with the highest posterior probability. Thus we are also able to visualize the spatial distribution of cell topics in the images.

#### Model Selection

The likelihood of the topic model is intractable to compute in general, but we can approximate the model log-likelihood in terms of model parameters {***β***_*k*_} and {***θ***_*d*_} [37]. With the law of total probabilities, we take into account uncertainties both in cells’ region and topic assignment, then the log-likelihood of the spatial topic model can be presented as

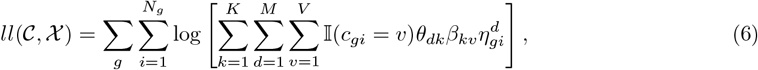

where 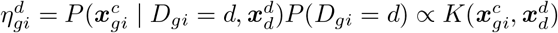.

We use the Deviance Information Criterion (DIC) [38] to select the number of topics, a generalization of the Akaike Information Criterion (AIC) in Bayesian model selection:

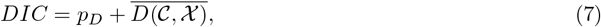

where the Deviance is defined as *D*(𝒞, *𝒳*) = −2*ll*(𝒞, *𝒳*) and 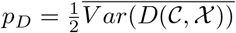.

DIC requires calculating the log-likelihood for every posterior sample, which is time-consuming. To determine the optimum number of topics, we run *SpatialTopic* with a varied number of topics (2-9 in practice) and collect a few posterior samples (such as the first 20 posterior samples) after convergence (with `trace=1`). The number of topics was selected based on DIC (7). Otherwise, we only output the deviance and the log-likelihood of the final posterior sample (default: `trace=0`).

#### Comparing to other methods

We compared the performance of *SpatialTopic* with five other niche analysis methods: spatial-LDA, Seurat-v5, UTAG, CytoCommunity, and BankSY. For BankSY and UTAG, we used protein or gene expression data and cell spatial coordinates as inputs, while the other methods used existing cell-type annotations and cell spatial coordinates. We followed the pre-processing procedures and parameters described in the original papers and tutorials for each method, with some hyperparameters slightly adjusted for computational efficiency on large datasets or when clear guidelines for tuning parameters were available. Details of these adjustments and the rationale for not using the default settings are described in this section.

All methods were initially run using R Studio (for R-based methods) or Jupyter Lab (for Python-based methods) on a standard MacBook Air (M2, 2022). If a method could not be run on a standard Mac due to memory constraints, we used our high-performance computing server with a single-core CPU and 200GB of assigned memory. For the Nanostring CosMx NSCLC dataset, both CytoCommunity and UTAG were run on the server due to high memory usage. Additionally, for the CODEX mouse spleen dataset, UTAG can be run on the Mac only without the default parallel mode due to memory constraints.

#### *SpatialTopic* (v1.1.0)

We ran *SpatialTopic* with region radius = 400, 150, 300 for the NSCLC, the mouse spleen, and the melanoma datasets, respectively, allowing around 100 cells per region on average during initialization, which is necessary for accurately estimating the topic-region distribution. We chose length-scale sigma = 20 for the mouse spleen dataset and used the default parameters for the NSCLC dataset. Posterior samples were collected after the convergence of the Gibbs sampling chain, with a burnin period of 2000 iterations for the NSCLC dataset and 1500 iterations for the mouse spleen dataset. For the Melanoma dataset, *SpatialTopic* was run with a burn-in period of 2000 iterations. For the healthy lung dataset with 26 small ROIs, *SpatialTopic* was run with sigma = 5 and radius = 60 to identify the complex local structures. In addition, we increase the number of initializations to 200 times to increase the robustness of identifying consensus patterns across ROIs while increasing the running time.

#### Seurat-v5 (v5.0.2)

We used the default niche analysis in Seurat v5, specifically the BuildNicheAssay() function in the Seurat R package. Seurat v5 employs k-means clustering to group cell neighborhood features, which are derived from the shared-nearest-neighbor graph (default neighbors.k = 30), a variant of the k-nearest-neighbor graph, as part of its image-based spatial data analysis pipeline. We ran BuildNicheAssay() with all default parameters except for the NSCLC datasets, for which we set neighbors.k = 100. We found that increasing neighbors.k from 10 to 100 (testing neighbors.k = 10, 30, 50, 100) significantly improved the algorithm’s performance on this dataset, with results presented in Figure S1.

#### Spatial-LDA (v0.1.3)

When working on mouse spleen datasets, we used the same parameters as the authors used in the original methodology paper, though we now use neighborhoods of all cells as the input, not only B cells. For the NSCLC datasets, we also use neighborhoods of all cells as the input but set radius = 400 to extract neighborhood cell type compositions. To reduce the computational complexity for both datasets, we set the threshold = 0.01 for ADMM Primal-Dual optimizer. Finally, we output the topic weights for every cell and assign every cell to a topic with the maximal weight.

#### CytoCommunity (Github version obtained on 2024 February)

CytoCommunity (unsupervised version) was run on a CPU with 200GB of assigned memory and evaluated only on the NSCLC dataset due to its demand for large-memory GPU resources and the unsupervised version’s inability to learn Tissue Cell Neighborhoods (TCNs) across multiple images (TCNs learned from individual images are not comparable). We set KNN-K = 300 for 0.1M cells, as suggested in the original paper. For large image data, the second step of CytoCommunity is time-consuming when trained on a CPU. Therefore, we greatly reduced num RUN to 10 and Num Epoch to 100 per run while ensuring the final loss was less than -0.2 for each run. Other parameters were set to their defaults.

#### UTAG (v0.1.1)

UTAG was primarily developed for protein expression data with limited marker channels. For the Nanostring CosMx NSCLC datasets with 960 genes, we used typical pre-processing steps suggested by Scanpy (v1.9.8) for analyzing scRNA-seq datasets. These steps included filtering low-prevalence genes, log transformation, and retaining only highly variable genes. We then performed z-score normalization, truncated at 10 standard deviations, followed by PCA. Only the top 50 principal components were used as input for UTAG. UTAG was run under multiple clustering resolutions [0.05, 0.1, 0.3, 0.5] and mix dist = 60, with an image resolution of 0.18 microns per pixel, since the authors suggested setting mix dist between 10 and 20 microns in the user manual. For the CODEX mouse spleen dataset (with intensity values already transformed), we performed z-score normalization truncated at 10 standard deviations, followed by Combat batch correction [31] and a second z-score normalization truncated at 10 standard deviations, a similar procedure as introduced in the UTAG paper for preprocessing IMC data [13]. We also set mix dist = 60, with an imaging resolution of 0.188 microns per pixel.

#### BankSY (v0.99.9)

In contrast to UTAG, BankSY is specifically designed to analyze spatial transcriptomics datasets. We ran BankSY with lambda = 0.8 to identify spatial domains, as recommended, with other parameters set to default, as described in the GitHub tutorial. For the NSCLC dataset, we followed the same pre-processing procedures outlined in the domain analysis tutorial, using k geom = 30, npcs = 50, and clustering resolutions of 0.1, 0.2, 0.3, and 0.5. For the mouse spleen datasets, we used the same input as UTAG, after batch correction and normalization. We followed the tutorial for multi-sample analysis, running the results under npcs = 30 since the dataset has only 30 markers.

### Data Preprocessing

#### Nanostring CosMx Human NSCLC

**s**The Nanostring CosMx NSCLC dataset is available on the Nanostring Website (https://nanostring.com/products/cosmx-spatial-molecularimager/ffpe-dataset/nsclc-ffpe-dataset/). For our analysis, we selected Lung5-1 sample and annotated about 0.1M cells into 38 cell types using Azimuth [23] with a human lung reference v1.0 (https://azimuth.hubmapconsortium.org/references/). We used the same cell annotations from the Seurat image analysis pipeline tutorial (https://satijalab.org/seurat/articles/seurat5spatialvignette2.html). Since healthy lung tissue was used as the reference, the ‘basal’ cells were re-labeled as tumor cells since they are the most closed cell type. We checked that the tumor locations indicated by the reference-based cell annotations are generally consistent with the tumor region labeled by the Nanostring company.

#### CODEX Mouse Spleen

We used the cell type annotation, marker expression level, and imaging coordinates from the original paper [2]. The image dataset can be downloaded from https://data.mendeley.com/datasets/zjnpwh8m5b/1. For cell coordinates, we only use the X and Y axes of the samples, ignoring Z axis. However, the result is similar when considering all three dimensions.

#### IMC Healthy Lung

We used the cell type annotation, marker expression level, cell imaging coordinates, and cell UTAG domain labels in the original paper [13]. This image dataset can be downloaded from https://zenodo.org/records/6376767.

#### mIF Melanoma

This is one of the whole-slide images from our internal mIF melanoma tissue samples [27]. Those whole tissue sections were stained using Ultivue UltiMapper I/O Immuno8 Kit (Cambridge, MA, USA) containing CD8, PD-1, PD-L1, CD68, CD3, CD20, FoxP3, and pancytokeratin + SOX10 (panCK-SOX10) followed by opal tyramide staining containing TCF1/7, TOX, Ki67, LAG-3. The whole imaging preprocessing pipeline has been previously described [27]. Here, we used only the cell phenotypes (classified based on marker expression of CD8, panCK-SOX10, CD68, CD3, CD20, FoxP3) and cell locations as the input of *SpatialTopic*.

### Simulation

We tested methods on simulated datasets of different scales to benchmark the scalability of *SpatialTopic* with an increasing number of cells in images. We randomly sampled 10k, 40k, 90k, 160k, and 250k pixels from an image, similar to the simulation method described in [21], to represent cell locations. We did not simulate gene expression levels for every individual cell. Instead, for each domain, we randomly sampled cells with domain-specific cell type distributions, with parameters simulated from *Dirichlet*(1, 1, 1, 1, 1), anticipating five distinct cell types per domain. Five unique datasets were generated for each simulation scenario. We also scaled the X and Y axes to maintain consistent cell densities across all simulation scenarios.

## Acknowledgements

We would like to thank computational support from MSK-MIND. This work is supported in part by the MSKCC Society, the V Foundation, the Parker Institute for Cancer Immunotherapy, NIH P30 CA008748, NIH R01 CA276286, and the MSK-MIND consortium.

## Data availability

All public datasets we used in the study can be downloaded online, with analysis details described in the Method section. Our in-house Melanoma dataset will be made public available with the analysis paper [27].

## Code availability

The R package is available on Github (https://github.com/xiyupeng/SpaTopic/) with a tutorial (https://xiyupeng.github.io/SpaTopic/). The R package is also available on CRAN (https://cloud.r-project.org/package=SpaTopic). The first version of the R package was officially released on CRAN on Jan 17, 2024.

## Author contribution

X.P. contributed to the original draft, developed the statistical model, and wrote the software. X.P., J.W.S., R.S., and K.S.P. developed the initial study concept. X.P., R.S., K.S.P., and J.L. developed the algorithm. X.P., C.E.K contributed to the R package. X.P. J.W.S., C.E.K, F.E., M.Y., M.B. analyzed the data. R.S., K.S.P., M.A.P, and M.K.C. oversaw all data generation and analysis. X.P., J.W.S., F.E., J.L., M.B., R.S., and K.S.P. edited the manuscript. All authors reviewed and approved the final manuscript.

## References

[1] Keren, L. et al. A Structured Tumor-Immune Microenvironment in Triple Negative Breast Cancer Revealed by Multiplexed Ion Beam Imaging. Cell 174, 1373–1387.e19 (2018).

[2] Goltsev, Y. et al. Deep Profiling of Mouse Splenic Architecture with CODEX Multiplexed Imaging. Cell 174, 968–981.e15 (2018).

[3] Ko, J. et al. Spatiotemporal multiplexed immunofluorescence imaging of living cells and tissues with bioorthogonal cycling of fluorescent probes. Nature Biotechnology 40, 1654–1662 (2022).

[4] Hoch, T. et al. Multiplexed imaging mass cytometry of the chemokine milieus in melanoma characterizes features of the response to immunotherapy. Science Immunology 7, eabk1692 (2022).

[5] Moldoveanu, D. et al. Spatially mapping the immune landscape of melanoma using imaging mass cytometry. Science Immunology 7, eabi5072 (2022).

[6] Nirmal, A. J. et al. The Spatial Landscape of Progression and Immunoediting in Primary Melanoma at Single-Cell Resolution. Cancer Discovery 12, 1518–1541 (2022).

[7] McCaffrey, E. F. et al. The immunoregulatory landscape of human tuberculosis granulomas. Nature Immunology 23, 318–329 (2022).

[8] Helmink, B. A. et al. B cells and tertiary lymphoid structures promote immunotherapy response. Nature 577, 549–555 (2020).

[9] Mature tertiary lymphoid structures predict immune checkpoint inhibitor efficacy in solid tumors independently of PD-L1 expression. Nature Cancer 2, 794–802 (2021).

[10] Cabrita, R. et al. Tertiary lymphoid structures improve immunotherapy and survival in melanoma. Nature 577, 561–565 (2020).

[11] Fridman, W. H. et al. B cells and tertiary lymphoid structures as determinants of tumour immune contexture and clinical outcome. Nature Reviews Clinical Oncology 19, 441–457 (2022).

[12] Feng, Y. et al. Spatial analysis with SPIAT and spaSim to characterize and simulate tissue microenvironments. Nature Communications 14, 2697 (2023).

[13] Kim, J. et al. Unsupervised discovery of tissue architecture in multiplexed imaging. Nature Methods 19, 1653–1661 (2022).

[14] Chen, Z., Soifer, I., Hilton, H., Keren, L. & Jojic, V. Modeling multiplexed images with spatial-lda reveals novel tissue microenvironments. Journal of Computational Biology (2020).

[15] Patrick, E. et al. Spatial analysis for highly multiplexed imaging data to identify tissue microenvironments. Cytometry Part A 103, 593–599 (2023).

[16] Hao, Y. et al. Dictionary learning for integrative, multimodal and scalable single-cell analysis. Nature Biotechnology 42, 293–304 (2024).

[17] Singhal, V. et al. BANKSY unifies cell typing and tissue domain segmentation for scalable spatial omics data analysis. Nature Genetics 56, 431–441 (2024).

[18] Hu, Y. et al. Unsupervised and supervised discovery of tissue cellular neighborhoods from cell phenotypes. Nature Methods 21, 267–278 (2024).

[19] Li, Z. & Zhou, X. BASS: multi-scale and multi-sample analysis enables accurate cell type clustering and spatial domain detection in spatial transcriptomic studies. Genome Biology 23, 1–35 (2022).

[20] Chidester, B., Zhou, T., Alam, S. & Ma, J. SpiceMix enables integrative single-cell spatial modeling of cell identity. Nature Genetics 55, 78–88 (2023).

[21] Shang, L. & Zhou, X. Spatially aware dimension reduction for spatial transcriptomics. Nature Communications 13, 1–22 (2022).

[22] Wang, X. & Grimson, E. Platt, J., Koller, D., Singer, Y. & Roweis, S. (eds) Spatial latent dirichlet allocation. (eds Platt, J., Koller, D., Singer, Y. & Roweis, S.) Advances in Neural Information Processing Systems, Vol. 20 (Curran Associates, Inc., 2007).

[23] Hao, Y. et al. Integrated analysis of multimodal single-cell data. Cell 184, 3573–3587 (2021).

[24] Rakaee, M. et al. Tertiary lymphoid structure score: a promising approach to refine the TNM staging in resected non-small cell lung cancer. British Journal of Cancer 124, 1680–1689 (2021).

[25] Aran, D. et al. Reference-based analysis of lung single-cell sequencing reveals a transitional profibrotic macrophage. Nature Immunology 20, 163–172 (2019).

[26] Meng, G. et al. imply: improving cell-type deconvolution accuracy using personalized reference profiles. Genome Medicine 16, 65 (2024).

[27] Smithy, J. W. et al. Spatial assessment of stromal b cell aggregates predicts response to checkpoint inhibitors in unresectable melanoma. medRxiv (2024).

[28] Peng, X. et al. A topic modeling approach reveals the dynamic T cell composition of peripheral blood during cancer immunotherapy. Cell Reports Methods 3, 100546 (2023).

[29] Roberts, M. E., Stewart, B. M. & Tingley, D. stm: An R Package for Structural Topic Models. Journal of Statistical Software 91, 1–40 (2019).

[30] Yosofvand, M. et al. Spatial immunophenotyping from whole-slide multiplexed tissue imaging using convolutional neural networks. bioRxiv (2024).

[31] Johnson, W. E., Li, C. & Rabinovic, A. Adjusting batch effects in microarray expression data using empirical Bayes methods. Biostatistics 8, 118–127 (2006).

[32] Kuett, L. et al. Three-dimensional imaging mass cytometry for highly multiplexed molecular and cellular mapping of tissues and the tumor microenvironment. Nature Cancer 3, 122–133 (2022).

[33] Datta, A., Banerjee, S., Finley, A. O. & Gelfand, A. E. Hierarchical nearest-neighbor gaussian process models for large geostatistical datasets. Journal of the American Statistical Association 111, 800–812 (2016).

[34] Weber, L. M., Saha, A., Datta, A., Hansen, K. D. & Hicks, S. C. nnSVG for the scalable identification of spatially variable genes using nearest-neighbor Gaussian processes. Nature Communications 14, 4059 (2023).

[35] Griffiths, T. L. & Steyvers, M. Finding scientific topics. Proceedings of the National Academy of Sciences 101, 5228–5235 (2004).

[36] Heinrich, G. Parameter estimation for text analysis (2009).

[37] Newman, D., Asuncion, A., Smyth, P. & Welling, M. Distributed algorithms for topic models. The Journal of Machine Learning Research 10, 1801–1828 (2009).

[38] Gelman, A., Carlin, J. B., Stern, H. S. & Rubin, D. B. Bayesian Data Analysis: Second Edition Texts in Statistical Science (CRC Press, 2004).

