## Supplemental Tables and Figures for "Decoding Spatial Tissue Architecture: A Scalable Bayesian Topic Model for Multiplexed Imaging Analysis"

### Supplementary Tables and Figures

Table S1: Real datasets used for benchmarking and illustration

| Datasets | Platform | Tissue | N_images | Reference |
| --- | --- | --- | --- | --- |
| Nanostring CosMx NSCLC | Nanostring CosMx | Human non-small cell lung cancer tissue | 1 | Nanostring Website |
| CODEX Mouse spleen | Co-detection by Indexing (CODEX) | Mouse spleen | 9 | Goltsev et.al.[1] |
| MIF Melanoma | Multiplexed immunofluorescence (MIF) | Melanoma soft tissue | 1 | Smithy et al.[2] |
| IMC Human healthy lung | Imaging Mass Cytometry (IMC) | Human healthy lung tissue | 26 | Kim et.al.[3] |

Table S2: Summary of methods compared in the benchmarking

| Method | Model | Input | Across slides? | Availability | Reference | Comments |
|---|---|---|---|---|---|---|
| Spatial-Topic | LDA with a flexible document design | Cell type, cell location | Yes | R package | This paper | |
| Spatial-LDA (2020) | Counting neighbors within a certain radius + LDA with a spatial prior | Cell type, cell location | Yes | Python library | Chen et. al.[4] | |
| Seurat v5 (2024) | SNN + Clustering (kmeans) | Cell type, cell location | Yes | R package | Hao et.al.[5] | |
| UTGA (2022) | KNN (average marker expression) + Clustering | Cell features (gene, morphology), cell location | Yes | Python library | Kim et.al.[3] | |
| CytoCommunity (2024) | Graph Neural Network+ majority voting | Cell type, cell location | Only supervised mode | Python library | Hu et.al.[6] | The TCNs learned across slides by unsupervised mode are not comparable. |
| BankSY (2024) | Neighborhood feature engineering (KNN) + PCA + Clustering | Cell features, cell location | Yes | R package | Singhal et.al.[7] | |

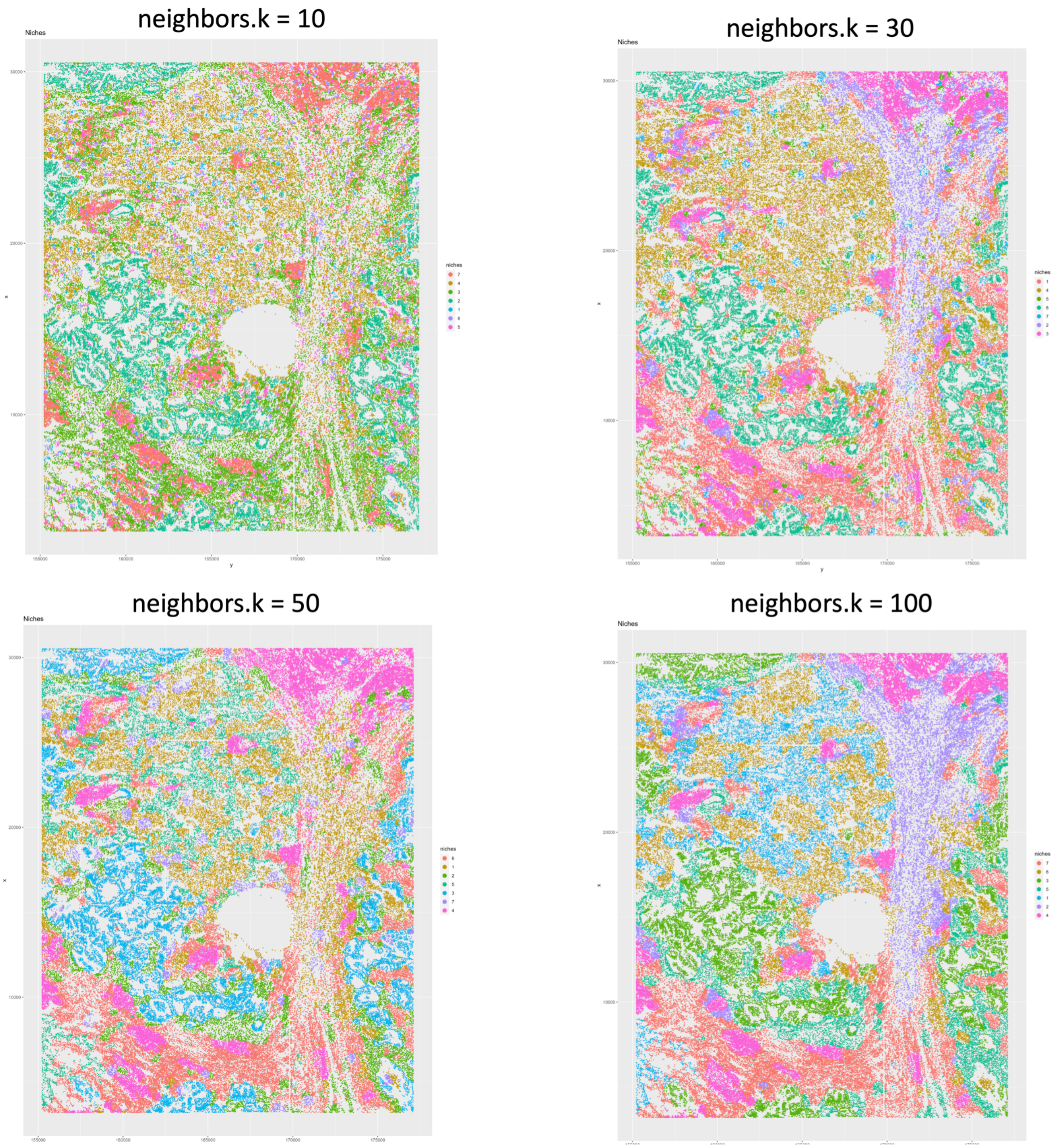


Figure S1. The effects of neighbors.k (number of neighbors selected when constructing the KNN graph) when applying the function BuildNicheAssay() of Seurat v5 to the CosMx NSCLC dataset (sample Lung5-1).

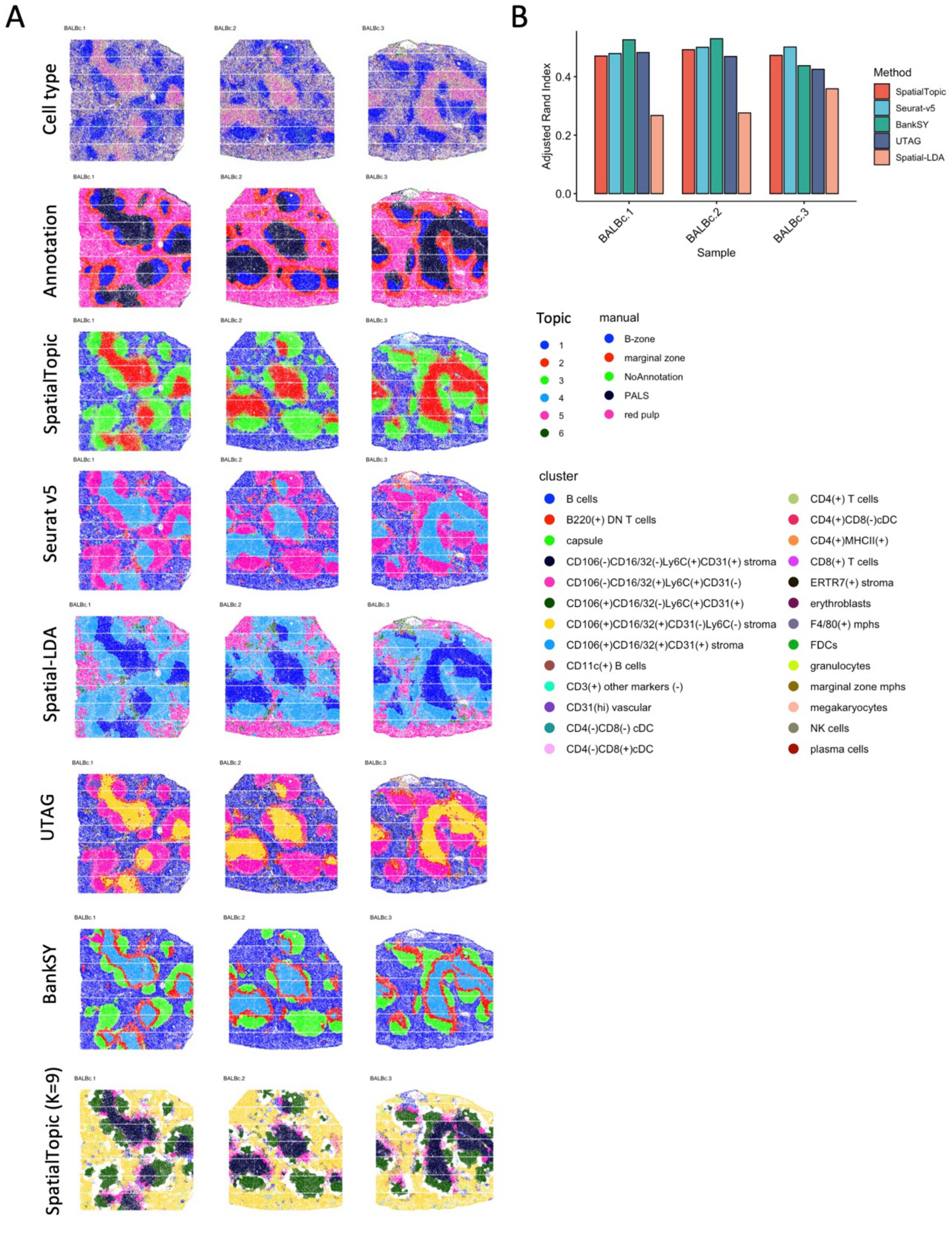


Figure S2: Benchmarking *SpatialTopic* compared to Seurat v5, Spatial-LDA, UTAG, and BankSY on the CODEX mouse spleen dataset.

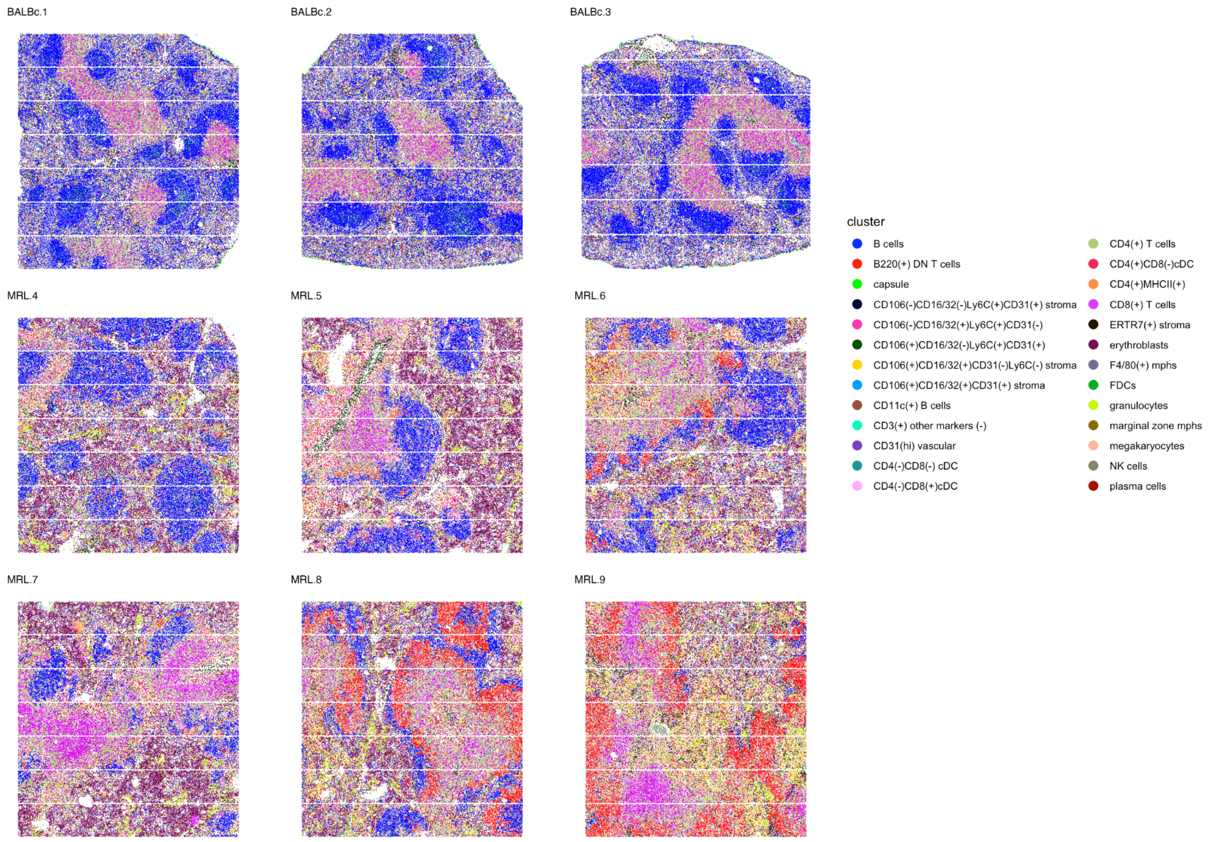


Figure S3: 27 major cell types identified across nine images in the CODEX mouse spleen dataset

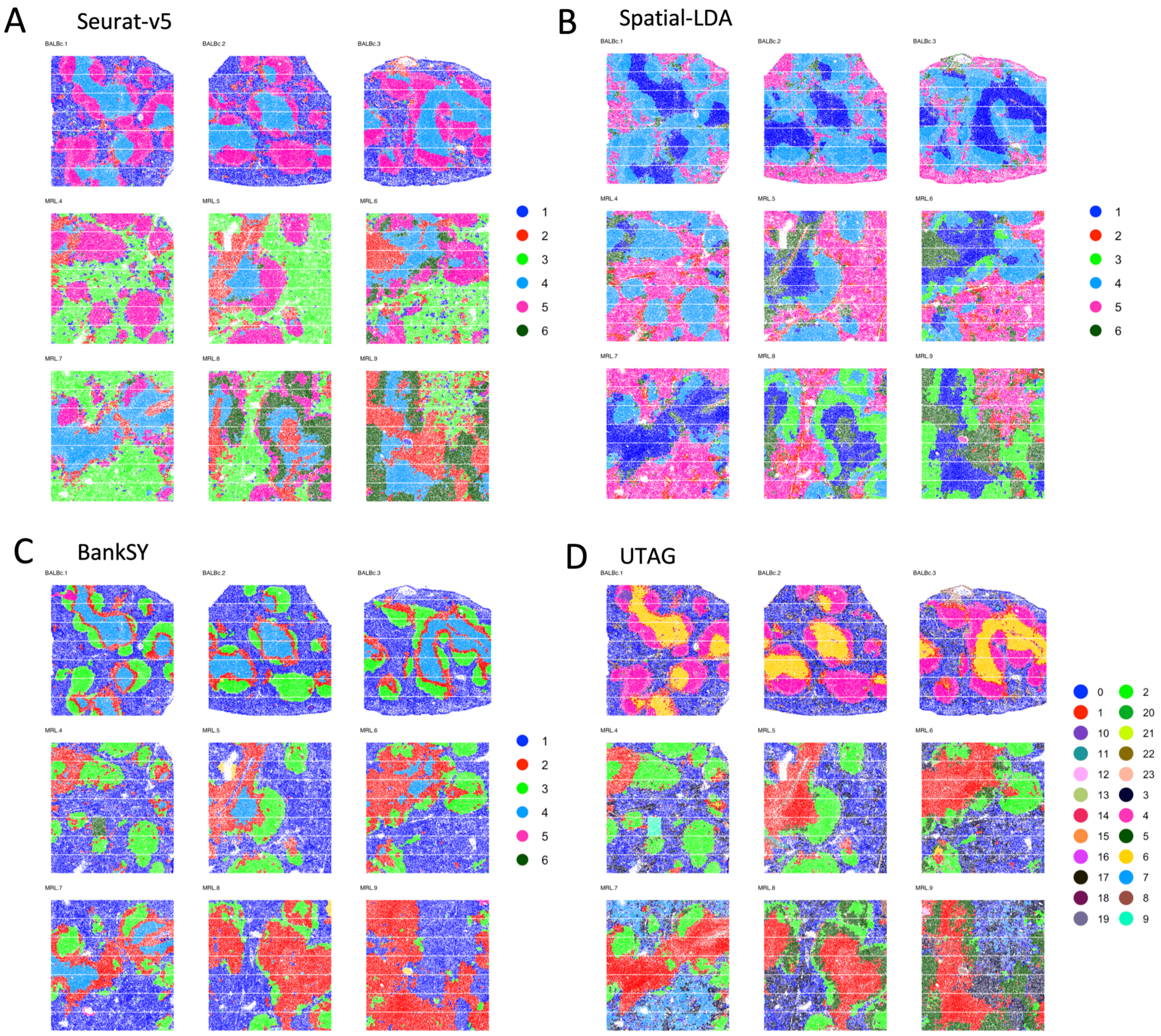


Figure S4. Benchmarking Seurat-v5, Spatial-LDA, BankSY, and UTAG on nine images of the CODEX mouse spleen dataset.

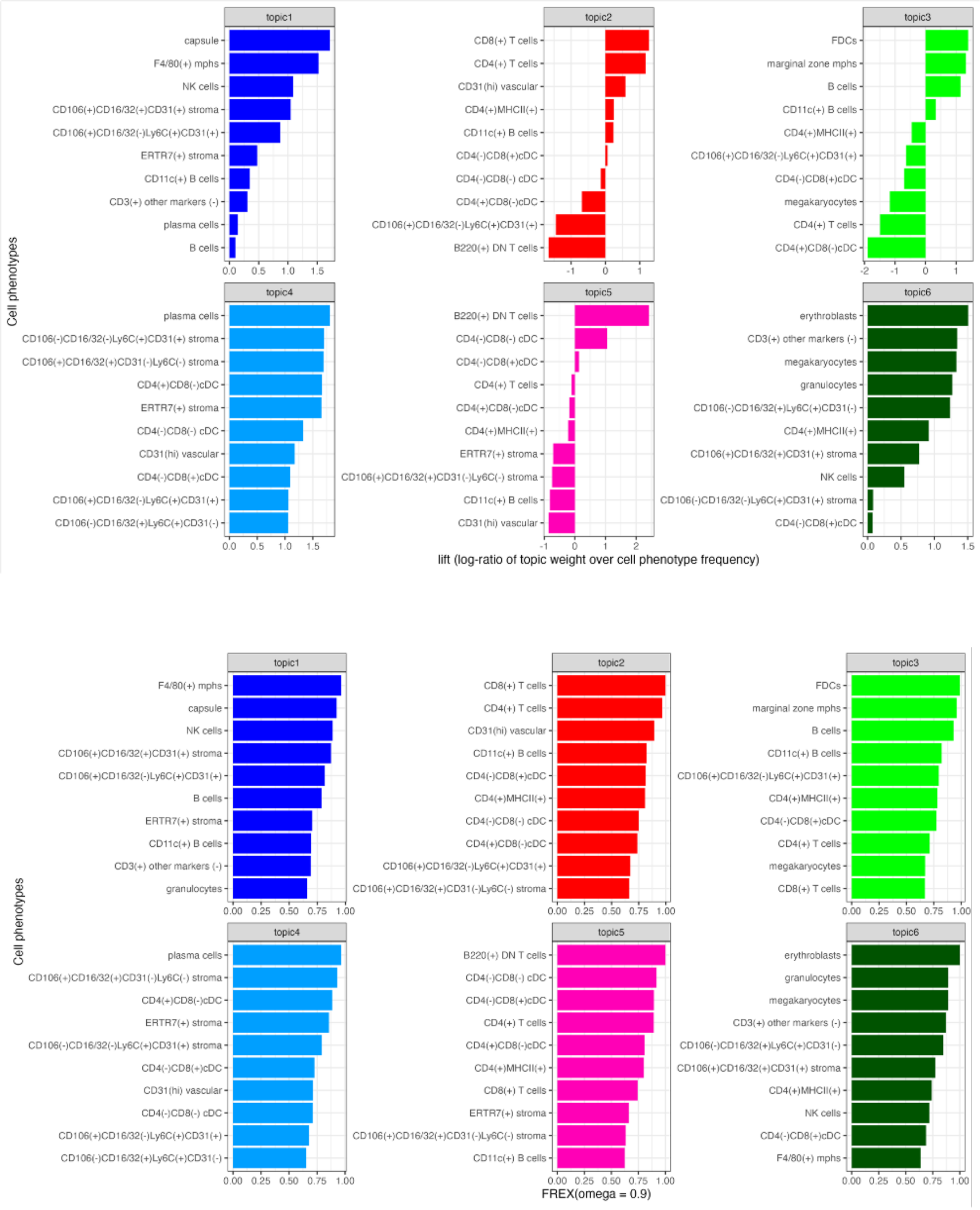


Figure S5. Topic-specific cell types identified by Lift metrics (upper) and FREX with omega = 0.9 (lower). For each topic, we select the top 3 cell types with the highest FREX (omega = 0.9), except for a few cases: For topic 4, we select ERTR7(+) stroma instead of plasma cells since plasma cells have very low abundance in the topic (See Figure 5C); We also drop CD4(-)CD8(+)cDC due to low lift.

### References


1. Goltsev, Y. *et al.* Deep Profiling of Mouse Splenic Architecture with CODEX Multiplexed Imaging. *Cell* **174**, 968-981.e15 (2018).
2. Smithy, J. W. *et al.* Spatial assessment of stromal B cell aggregates predicts response to checkpoint inhibitors in unresectable melanoma. *medRxiv* 2024.08.09.24311758 (2024) doi:10.1101/2024.08.09.24311758.
3. Kim, J. *et al.* Unsupervised discovery of tissue architecture in multiplexed imaging. *Nat. Methods* **19**, 1653–1661 (2022).
4. Chen, Z., Soifer, I., Hilton, H., Keren, L. & Jojic, V. Modeling Multiplexed Images with Spatial-LDA Reveals Novel Tissue Microenvironments. *J. Comput. Biol.* **27**, 1204–1218 (2020).
5. Hao, Y. *et al.* Dictionary learning for integrative, multimodal and scalable single-cell analysis. *Nat. Biotechnol.* **42**, 293–304 (2024).
6. Hu, Y. *et al.* Unsupervised and supervised discovery of tissue cellular neighborhoods from cell phenotypes. *Nat. Methods* **21**, 267–278 (2024).
7. Singhal, V. *et al.* BANKSY unifies cell typing and tissue domain segmentation for scalable spatial omics data analysis. *Nat. Genet.* **56**, 431–441 (2024).
